## Supplemental Figures for "Unveiling Liquid-Crystalline Lipids in the Urothelial Membrane through Cryo-EM"

**Supplemental Materials**

**Supplemental Figures 1-4**

**Supplemental Table 1**

**Supplemental Spread sheet 1**

**Supplemental Movie 1**

**Supplemental Data 1**

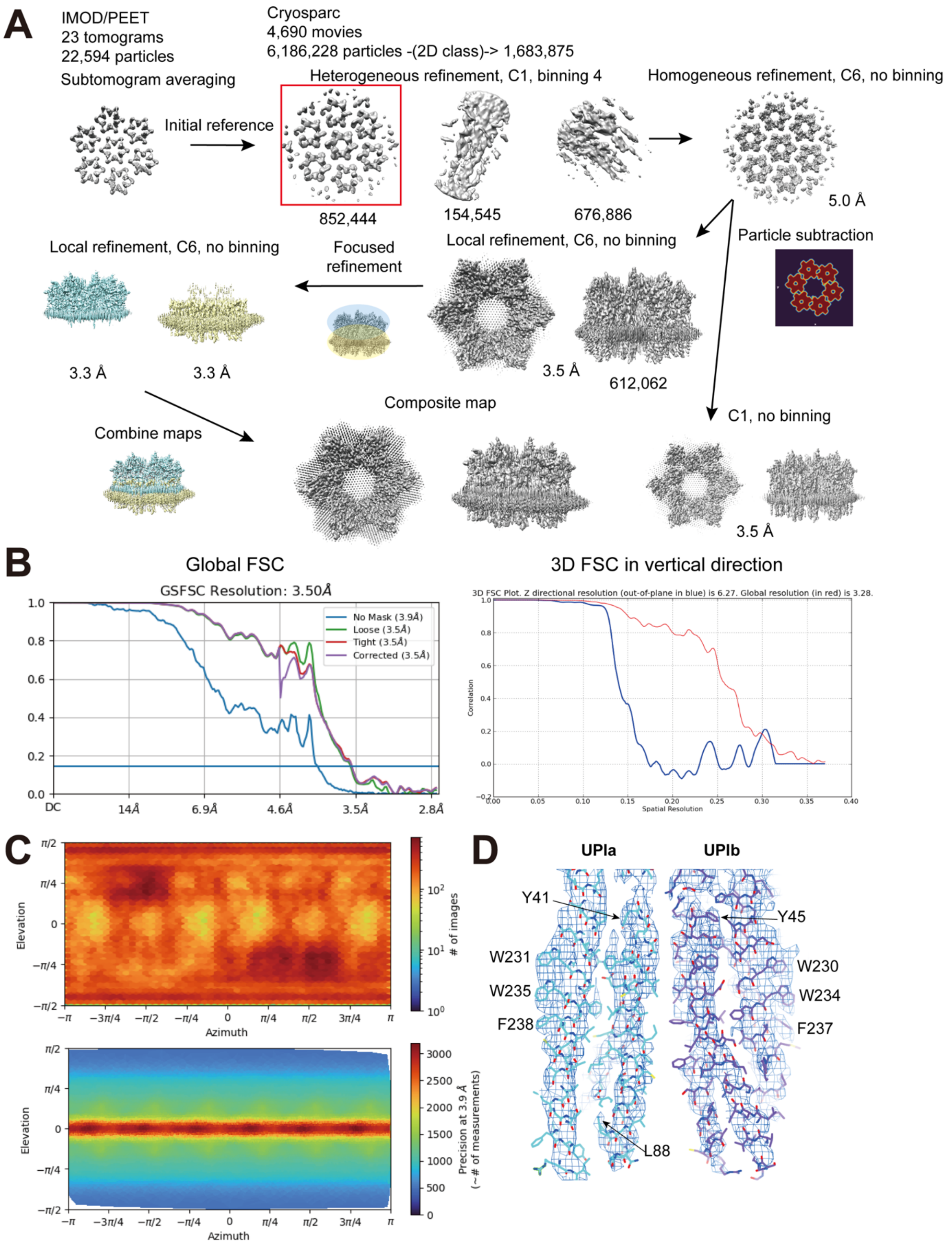

**Supplementary Figure 1**: Cryo-electron microscopy of the AUM. (A) Summary of the reconstruction procedure. The hexagonal lipid array was observed without applying C6 symmetry. (B) Resolution estimations using Fourier shell correlation. The cut-off value is

0.143. The 3D FSC in the vertical direction was shown on the right. (C) The viewing direction distribution (top) and the posterior precision directional distribution (bottom). Although the distribution of the views covers nearly entire directions, the precision of the side views deteriorates due to the overlapping molecules. (D) Transmembrane helices of UPIa and UPIb were shown along with fitted models to present the quality of the map.

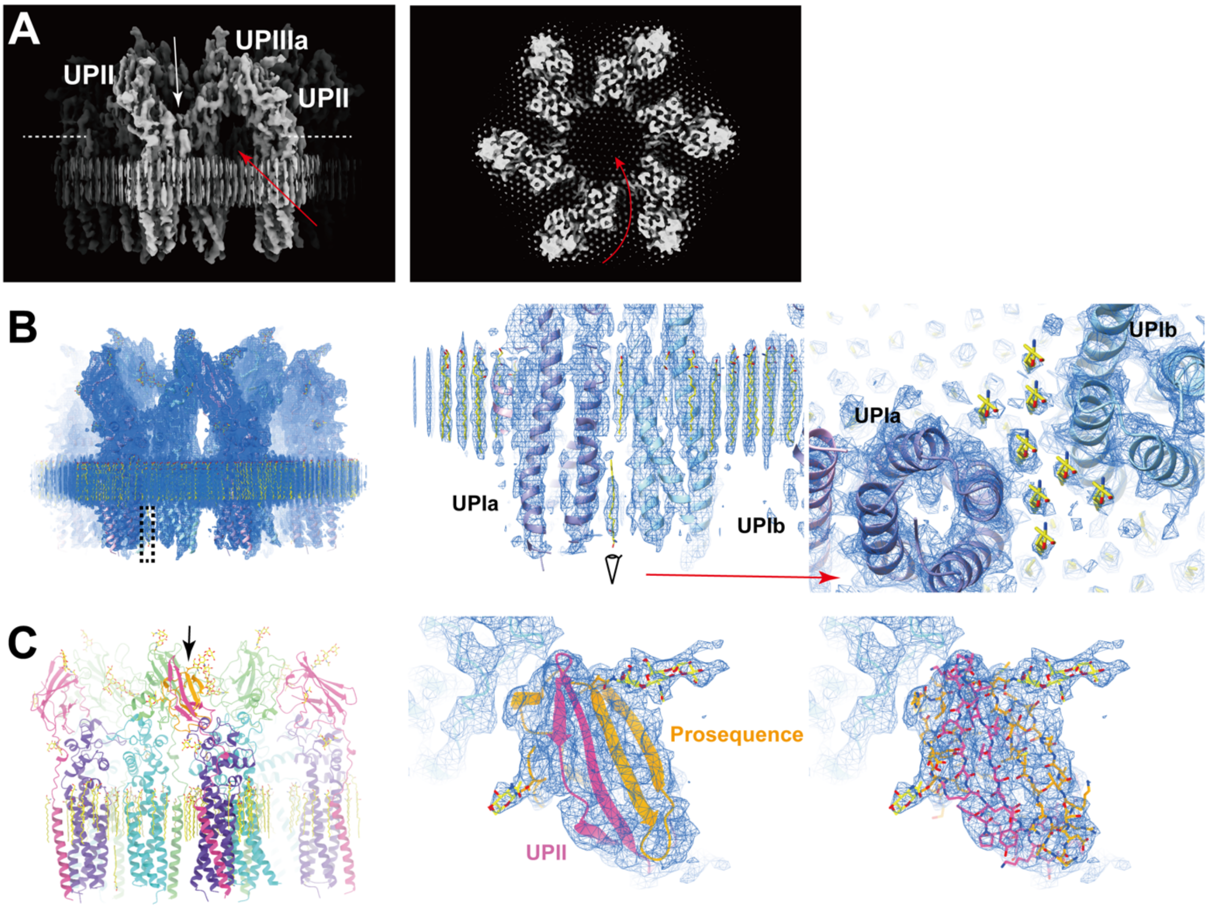

**Supplementary Figure 2**: Additional structural features. (A) Y-shaped conformation of the uroplakin heterotetramer. The white and red arrows indicate the groove between UPII and UPIIIa, and the channel beneath the II-IIIa arch, respectively. The broken lines indicate the position of the cross-section on the right panal. (B) Weak signals of lipids of the inner leaflet (arrow). The view from the cytoplasmic side (right) shows the hexagonal alignment of the inner lipids (yellow). (C) The prosequence of UPII (arrow, models in orange) forms an anti-parallel beta-sheet structure. The electron densities (mesh) validate the assignment of the domain (middle and right).

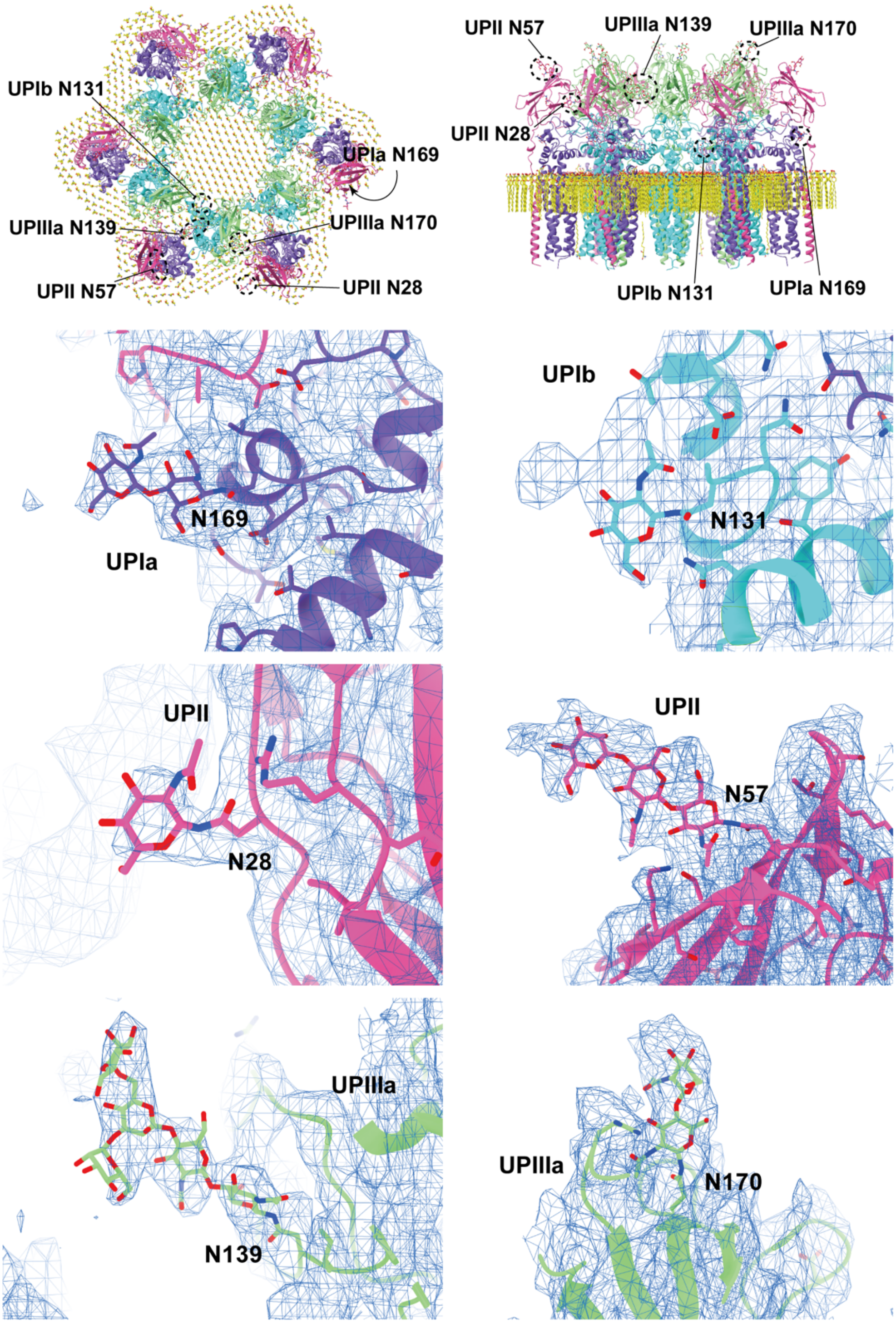

**Supplementary Figure 3:** Glycosylation of the uroplakin subunits. Six N-glycosylation sites were visualized in the reconstruction. The large densities of the carbohydrate chains at Asn139 of UPIIIa may interact with the N-terminal loop of the adjacent IIIa. Asn19 of UPII, which was also predicted to be glycosylated (Hu et al., 2005), was not observed in our map, probably due to the cleavage of the N-terminal signal sequence.

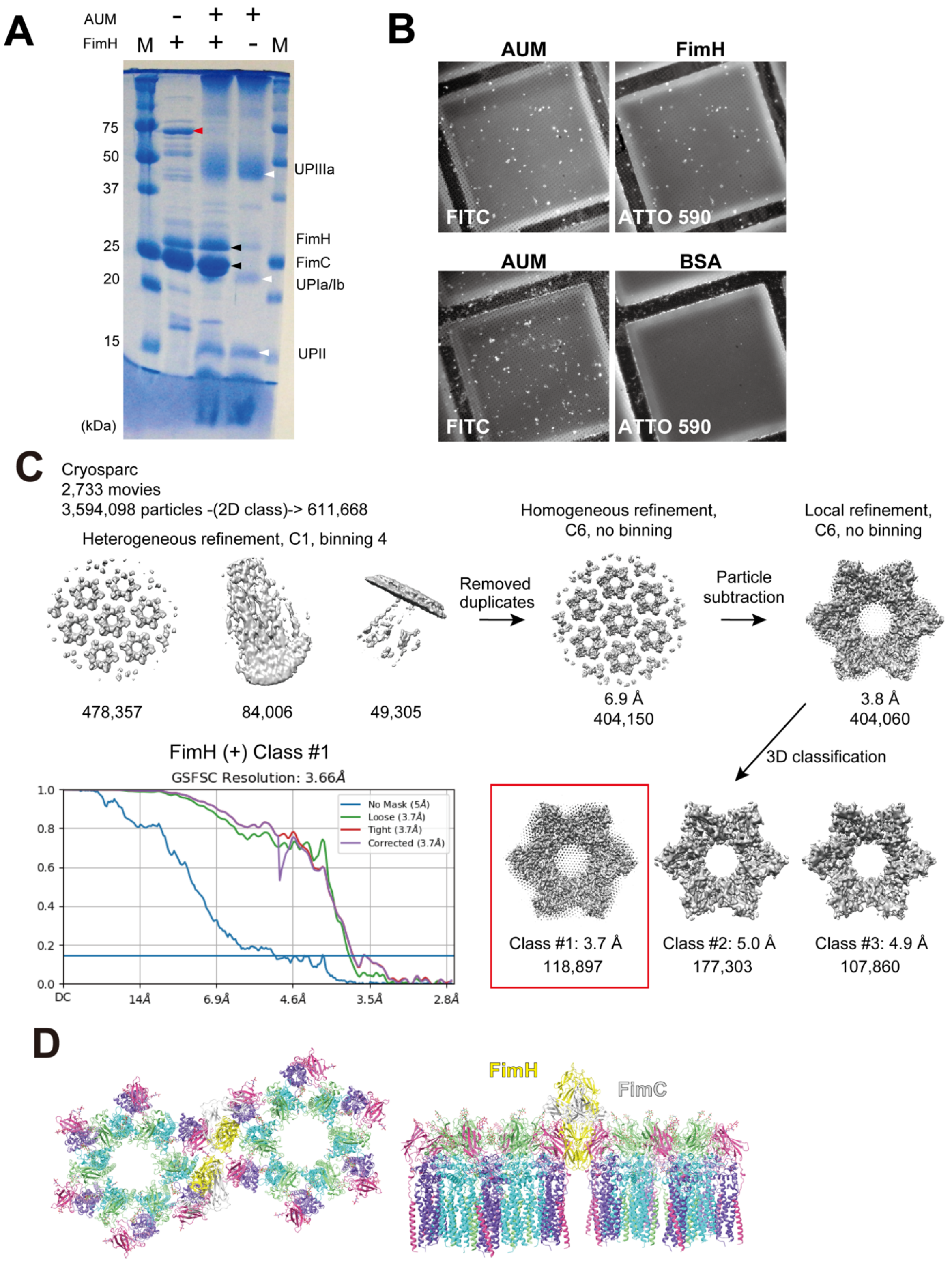

**Supplementary Figure 4**: **Binding of FimH to the AUM.** (A) SDS-PAGE of FimH-bound AUM. Purified FimH-FimC complexes were mixed with the AUM, and unbound molecules were removed through three rounds of centrifugation. Black arrowheads indicate FimH and

FimC; white arrowheads point to UP subunits; the red arrowhead highlights the *E. coli* heat shock protein 70, which was present as a contamination in the purified FimH-FimC sample. Notably, this protein does not bind to the AUM and is absent in the AUM mixed sample. The CBB staining of UP subunits appears weakened due to intensive glycosylation. (B) Fluorescence microscopy of frozen-thawed grids. FITC-labeled AUM mixed with ATTO 590-labeled FimH-FimC or ATTO 590-labeled BSA were loaded onto Quantifoil grids and frozen using liquid ethane. After thawing, the grids were imaged. The co-localization of FITC-labeled AUM and ATTO 590-labeled FimH-FimC signals confirms the presence of FimH/FimC on the AUM. In contrast, ATTO 590-labeled BSA did not attach to the AUM. (C) Summary of the reconstruction procedure. The Fourier shell correlation curve of class #1 is displayed. (D) Model of the uroplakin complex pair with bound FimH-FimC complexes (PDBID: 1QUN). The gap between two neighboring uroplakin complexes is so narrow that the mannose-binding domain of FimH barely fits.

**Supplementary Table 1: Summary of data collection and model validation**

| **Data collection parameters** | |
|---|---|
| Magnification | 64,000× |
| Pixel size (Å) | 1.35 |
| Voltage (keV) | 300 |
| | Tomography |
| Defocus range (µm) | 2.5-4.5 |
| Exposure time (sec/tilt) | 0.18 |
| Number of frames per tilt | 10 |

| | |
|---|---|
| Angular range (°) | ±60 |
| Increments (°) | 3 |
| Total dose ($e^{-}$/Å$^{2}$) | 50 |
| Number of tilt series recorded | 95 |
| Number of tilt series processed | 23 |
| Initial number of subtomograms | 83,109 |
| Final number of subtomograms | 22,594 |
| | SPA |
| Defocus range (µm) | 1.1-4.5 |
| Exposure time (sec) | 6.7 |
| Number of frames | 50 |
| Total dose ($e^{-}$/Å$^{2}$) | 50 |
| Tilt angles (°) | 0, 30 , 45, 55 |
| Ratio of tilted images | 1:1:2:2 for 0, 30, 45, 55° |
| Number of movies recorded | 4,690 (FimH-)/2,733 (FimH+) |
| Box size (pixels) | 512 |
| Initially picked particles | 12,764,463 (FimH-)/6,619,664 (FimH+) |
| Final particles | 609,567 (FimH-)/118,897 (FimH+) |
| **Model validation statistics** | |

| | |
|---|---|
| Initial model used | AlphaFold prediction |
| Bonds length RSMD (Å) | 0.023 |
| Bonds angles RSMD (°) | 2.466 |
| MolProbability score | 1.74 |
| Clash score | 2.55 |
| Rotamer outliers (%) | 1.49 |
| Ramachandran plot (%) | Favored: 89.25<br>Allowed: 10.75<br>Outliers: 0.00 |
| CaBLAM outliers (%) | 4.37 |
| B-factors (min/max/mean) | 59.32/134.09/93.14 |
| Map resolution estimates (Å) | 3.5 ($FSC_{half\text{-}map}$=0.143)<br>3.6 ($FSC_{model}$=0.143) |

**Supplemental spread sheet 1**: Numerical data used for generating boxplot in Figure 6.

**Supplemental Movie 1**: Movie representation of Figure 2A.

**Supplemental Data 1**: Maps and models used in this manuscript.

8JJ5.pdb: models of the uroplakin heterotetramer.

Associated_lipids.pdb: models of sphingosine/ceramides associated with the uroplakin complexes.

Other_lipids.mmcif: models of sphingosine surrounding the uroplakin complexes.

EMD-36340.mrc: electron density map of the uroplakin hexameric complex.

EMD-36340-DeepEMhancer_sharpened.mrc: sharpened map using DeepEMhancer tool of CryoSPARC.

C1map.mrc: electron density map without applying C6 symmetry.

FimHplus_Class1/2/3.mrc: FimH-bound structures.

Alphafold_prediction.pdb: the initial model predicted using Alphafold.

36340_8JJ5_ChimeraX_session.csx: session command file for visualization of maps and models using ChimeraX.
